# Not scene learning, but attentional processing is superior in team sport athletes and action video game players

**DOI:** 10.1101/353953

**Authors:** Anne Schmidt, Franziska Geringswald, Fariba Sharifian, Stefan Pollmann

**Affiliations:** Department of Experimental Psychology, Otto-von-Guericke University, 39106 Magdeburg, Germany; Transfer and Entrepreneur Centre, Otto-von-Guericke University, 39106 Magdeburg, Germany; Center for Behavioral Brain Sciences, 39106 Magdeburg, Germany

**Keywords:** visual search, attention, spatial memory, handball, team sports, action video game playing

## Abstract

We tested if high-level athletes or action video game players have superior context learning skills. Incidental context learning was tested in a spatial contextual cueing paradigm. We found comparable contextual cueing of visual search in repeated displays in high-level amateur handball players, dedicated action video game players and normal controls. In contrast, both handball players and action video game players showed faster search than controls, measured as search time per display item, independent of display repetition. Thus, our data do not indicate superior context learning skills in athletes or action video game players. Rather, both groups showed more efficient visual search in abstract displays that were not related to sport-specific situations.

---

Team sport athletes have shown superior performance in visuospatial attentional processing in a multitude of tasks (Mann, Williams, Ward, & Janelle, 2007). However, investigation with athletes often focus on sport-specific situations (Abernethy, 1991; Timmis, Turnmer, & van Paridon, 2014) making it difficult to infer general underlying processes. A recent study, however, found improved skills in athletes in a neutral, non-sport specific task with substantial visuospatial demands (Faubert, 2013). In this study, professional team sport players and high-level amateur players showed steeper learning curves in a repeatedly presented multiple object tracking (MOT) task (Pylyshyn, & Storm, 1988) than non-athletic controls. In this task, multiple dots out of a larger group are marked as targets at the beginning of a trial. Then the marking disappears, and all dots start to move on individually different random paths for several seconds. When the dot movement stops, the target dots need to be indicated. Their superior performance led to the claim that “professional athletes as a group have extraordinary skills for rapidly learning unpredictable, complex dynamic visual scenes that are void of any specific context” (Faubert, 2013, p. 3). However, the only learning that may influence performance in the MOT task is a procedural learning of the dynamic allocation of spatial attention (Alvarez, & Franconeri, 2007; Merkel, Hopf, & Schoenfeld, 2017) because the dot movements in the MOT-task are randomly generated, so there is no predictive information either within a trial or across different trials (“dynamic scenes“) that could be learned and used for memory-guided search. However, even though, to our knowledge, there is no empirical evidence for enhanced memory-guided attentional selection in team sport athletes, this is nevertheless an interesting question. It may well be that these athletes have superior capabilities to learn scenes and use scene memory for attentional guidance when the same or similar scenes are repeatedly encountered. Handball players, for example, need to move in a particular direction or pass the ball to a specific team member in a fraction of a second to be successful players. Furthermore, specific situations are repeatedly encountered during a game and may facilitate selection of the appropriate action. Thus, it could be that elite team sport athletes have extraordinary skills in learning spatial contexts and using context-knowledge for efficient attentional guidance in scenes that have been encountered before. If these skills - due to training or as a selection phenomenon (Kristjansson, 2013) - transfer to non-sport-specific situations, they would lead to benefits outside of sport and should be observed even in abstract (semantically meaningless) search tasks. This is what we investigated here in a group of high-level amateur handball players. We compared their performance to that of nonathletes on the one hand and to action video game players on the other hand. Action video game players were selected because enhanced attentional skills have been reported in this group, including improved visual control, greater attentional capacity, and better spatial allocation of attention (Green & Bavelier, 2003), enhanced target detection (Feng, Spence, & Pratt, 2007; Green & Bavelier, 2006a), and faster response selection (Castel, Pratt, & Drummond, 2005; Clark, Lanphear, & Riddick, 1987). For example, visual search was improved for action video game players relative to non-gamers (Bavelier, Green, Han, Renshaw, Merzenich, & Gentile, 2011; Buckley, Codina, Bhardwaj, & Pascalis, 2010; Chisholm, Hickey, Theeuwes, & Kingstone, 2010; Chisholm & Kingstone, 2012; Green & Bavelier, 2007; Hubert-Wallander, Green, Sugarman, & Bavelier, 2011).

However, conclusions regarding superior attentional skills in video gamers have also met scepticism. Some issues have been different motivational levels of expert and control groups, differential training histories and the typical gender imbalance of video game studies are discussed by Kristjánsson (2013). Moreover, due to sometimes inadequate control conditions claims of video game advantages should be taken as tentative (Boot et al., 2011).

Like playing of a team sport, action video game playing leads to complex sensory stimulation. In laboratory settings, action video game players showed faster visual search in symbolic, not game-related displays (Castel et al., 2005; Hubert-Wallander, Green, Sugarman, & Bavelier 2011; Wu, & Spence, 2013). An obvious difference is the complexity of visuo-motor demands that is much higher for handball players and may additionally support learning (Kramer, & Erickson, 2007).

A causal relationship between action video game experience and enhanced visual and cognitive performance (Green & Bavelier, 2003; Li, Polat, Makous, & Bavelier, 2009) and enhanced attentional processes and resource management has been postulated (Green & Bavelier, 2012). This, however, is still debated. Issues like gender imbalance, different motivational levels, differential training histories of testing groups, lack of comparable groups of video game studies and inadequate control conditions have been criticized (Boot et al., 2011, 2013; Kristjánsson, 2013).

To investigate the contribution of scene learning to attentional processing in these groups, we used the contextual cueing task (Chun, & Jiang, 1998). In this visual search paradigm, a target element has to be searched in a distractor-filled display.

A typical block of trials consisted of one half of displays that were repeatedly presented in subsequent blocks, while the other half of displays was always randomly generated. In numerous studies with this paradigm, it has been found that search becomes more efficient in repeated displays, even though participants are often unaware of these repetitions (Chun, 2000). This search time advantage for repeated versus novel displays indicates memory guided search. In addition to this specific scene learning effect, unspecific task learning effects can be assessed by the learning curves across novel and repeated displays. The nature of these learning effects can be broken down even further by assessing the slope and intercept of the search time function (response time as a function of the number of display elements; Kunar, Flusberg, Horowitz, & Wolfe, 2007). This analysis is based on a two-stage model of attentional processes, that are active during search and postselective processes that follow target selection. The slope indicates the increase of search times with increasing elements in the search display. It can be quantified as search time per item and thus gives an estimate of attentional processing speed. This should not be taken literally, shallower search slopes can reflect faster sequential search or more parallel search. For instance, the ability to process more items during a fixation (a larger attentional focus) would lead to shallower slopes. In the case of serial and parallel search alike, however, the search slope reflects the efficiency of the search. In contrast, the intercept of the search time function with the y-axis indicates the residual time needed for postselective processes that are independent of the number of display elements, in particular processes for preparing and executing the response.

In the contextual cueing experiment, participants had to search for a T-shape among arbitrary configurations of L-shaped distractors (Figure 1). Unbeknownst to the participants, one half of the configurations was repeatedly presented, so that search could be guided to the target location by incidentally learned configurations. Incidental learning was characterized by faster search in repeated configurations than novel configurations. Previous eye tracking studies have shown that search in repeated displays is characterized by a monotonic approach phase that typically begins after a few exploratory fixations. In this approach phase, each new fixation is nearer to the target than the previous one (Manginelli, and Pollmann, 2009; Tseng, and Li, 2004). This pattern suggests an ongoing process of matching aspects of the search display with the memory trace of previous displays held in working memory. Indeed, the search facilitation of repeated displays is dependent on visuospatial working memory, as demonstrated by the loss of the search facilitation when working memory is loaded by a secondary task (Manginelli, Geringswald, and Pollmann, 2012; Manginelli, Langer, Klose, and Pollmann, 2013).

**Figure 1:**
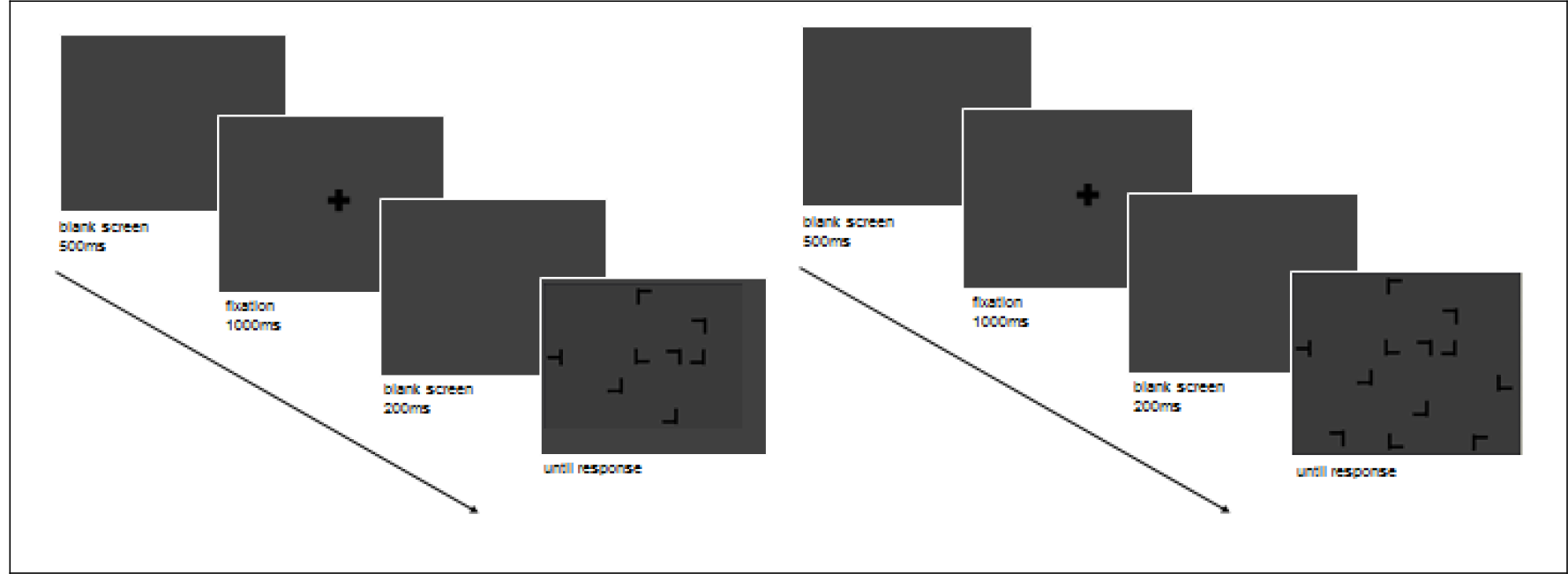
Schema of an experimental trial. Each trial consisted of a blank interval (500 ms) followed by the fixation cross (1000 ms) and a brief pause of 200 ms. Subsequently, participants were asked to search for a tilted target letter T among L-shaped distractors and the search display was presented until a response was made, followed by auditory feedback for the correctness of the response. Set size was varied between seven distractors (left) and eleven distractors (right) randomly within search blocks.

If athletes or action video game players had an improved capacity for learning visuospatial configurations and using them for memory-guided search, we expected an increased search facilitation (reduction of search times) in these groups relative to the control group in repeated compared to novel displays. Note that we tested these abilities in a setting that is not sport-specific, in order to see if athletes show generalized improvements of contextual cueing. If, alternatively, team sport athletes or action video game players have superior attentional capabilities independent of memory, they would be faster than controls in searching novel and repeated displays alike and, additionally, search slopes should be shallower than in normal controls. A third potential hypothesis would be improved response (including response preparatory) processes in team sport athletes and action video game players. This would lead to reduced intercept values of the search time function.

## Methods

### Participants

A total of 75 participants (control: n=25 (10 men, 15 women), mean age = 24.4 years, SD = 3.4; handball players: n=25 (15 men, 10 women), mean age = 21.5 years, SD = 7.1; action video game players: n=25 (20 men, 5 women); mean age = 25.2 years, SD = 7.1) were recruited from the University of Magdeburg and the Olympic Training Center Saxony-Anhalt / Sportclub Magdeburg. All subjects were students or had already completed an academic education. The handball players consisted of fifteen adult players (5 men, 10 women) from a 3rd league handball team and 10 junior players (10 men) playing in the A and B national youth league. None of the handball players was a regular video game player, as assessed by interview. Video game players and controls were recruited from the University of Magdeburg. Action video game players had to fulfill the following criteria: action video game players needed to play action video games (e.g. Call of Duty; Activision, Infinity Ward) for a minimum of five hours a week for at least one year. Participants without any team sport and little to no action video game experience (less than 1 hour per week) were classified as controls. Twenty-one participants already participated in previous contextual cueing experiments, however with different repeated displays (12 handball players, 9 video game players). All participants had normal or corrected-to-normal vision.

Informed written consent was acquired prior to the experiments. Subjects were remunerated with course credits or received a modest payment. Further, subjects were naive about the purpose of the experiment. The experiment was carried out in accordance with the declaration of Helsinki and was approved by the Ethics Committee of the University of Magdeburg.

### Apparatus & Stimuli

The experiment was run using the OpenGL-Psychophysics Toolbox extensions (Brainard, 1997; Pelli, 1997) in MATLAB (The MathWorks, Sherborn, MA). Stimuli were displayed by a projector on a back-projection screen (1170 mm (1024 pixels) wide and 850 mm (768 pixels); vertical refresh rate of 60 Hz). Participants viewed the stimuli from a distance of 126 cm (pixel size of 0.048° x 0.046°). Subjects completed the experiment individually in a dimly lit, sound-attenuated chamber. Search displays contained one target (90° or 270° rotated T) and 7 or 11 distractors (0°, 90°, 180°, 270° rotated L) with each item subtending 1.9° × 1.9°. An offset of 0.14° between the two segments of the L-shapes was chosen to increase search difficulty. The orientation of the target and the orientation of distractors were randomly chosen for each trial. A black cross (2.4° × 2.4°) at the center of the display was used as a fixation stimulus. Stimuli were black displayed on a gray background. The items were randomly positioned on four imaginary concentric circles with radii of 4°, 8°,12°, and, 16° each corresponding to 4, 12, 20, and 28 equidistant possible item locations. The visual angle of the search display on the projection screen extended an area of 49.8° x 37.3°.

Each trial started with a blank interval of 500 ms followed by the fixation cross for 1000 ms. After a brief pause of 200 ms, the search display appeared on the projection screen (Figure 1). Participants were asked to search for the target letter T among L-shaped distractors and further to specify as quickly and accurately as possible whether the stem of the T was pointing to the left or right by mouse button presses. They were further instructed not to apply active search strategies, because these strategies diminish contextual cueing (Lleras, & von Mühlenen, 2004). The search display remained on the screen until a response was made by the participants. Auditory feedback was provided for correct (a 500-Hz low-pitch tone) and incorrect answers (a 1500-Hz high-pitch tone). Blocks comprised 24 trials, 12 for each configuration type (repeated vs. novel). The positions of the items were balanced across quadrants and configuration type. Half of the trials had a set size of 8 items, whereas the other half of trials comprise 12 items, presented in randomized order (Figure 2).

**Figure 2:**
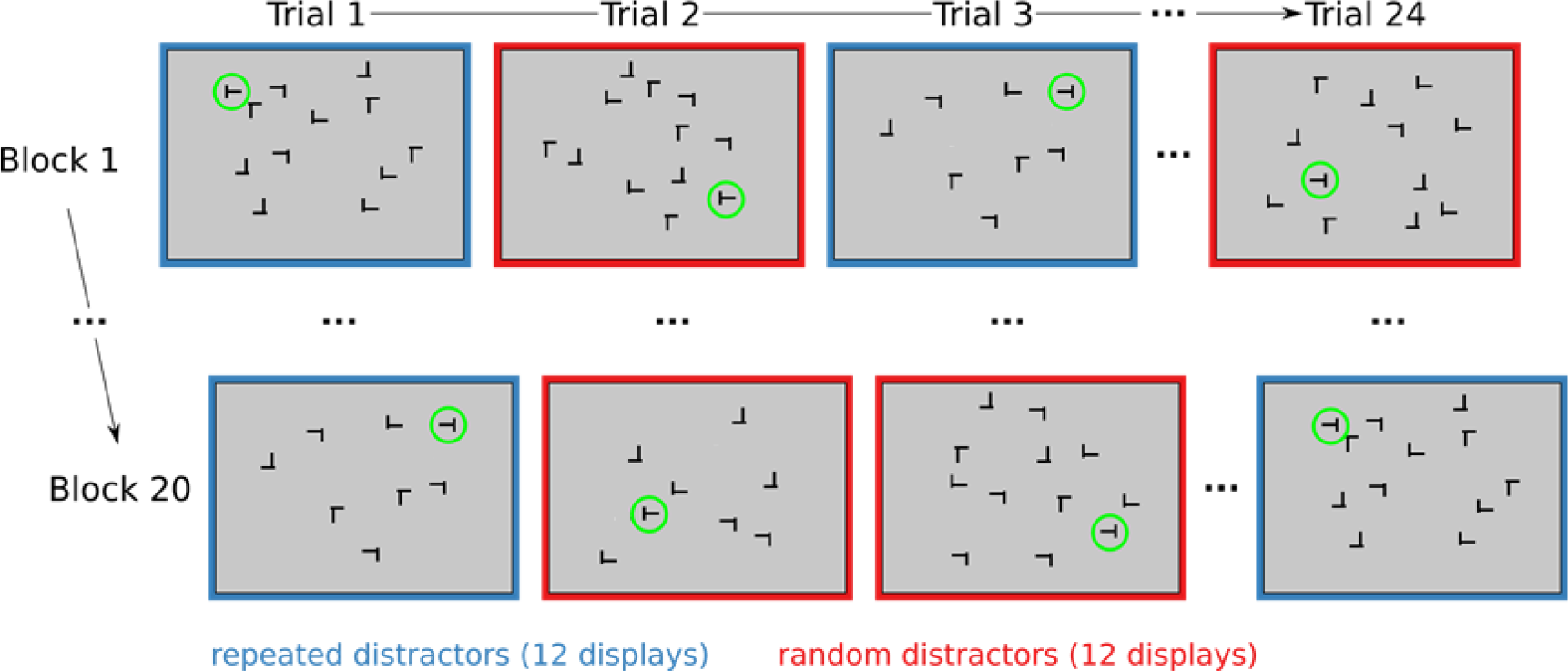
Experimental procedure. 20 blocks of 24 trials were presented. Blocks comprised 24 trials, 12 for each configuration type (repeated vs. novel). The red (novel displays) and blue (repeated displays) frames and the green circles indicating the targets are added for clarity, they were not visible during the experiment. The positions of the items were balanced across quadrants and configuration type. Half of the trials had a set size of 8 items, whereas the other half of trials comprise 12 items, presented in randomized order.

The experiment started with 24 randomly generated displays (which were not analyzed) to familiarize the participants with the task. Immediately afterwards 20 blocks of 24 trials were presented. Novel and repeated displays were created in the exact same way. The only difference was that in blocks 2-20, 12 of the displays of block 1 were repeated, while the remaining 12 were newly created for each block. The entire experiment lasted approximately 45 min. At the end of the search task, the participants performed a recognition test, to evaluate whether repeated displays were explicitly remembered. The recognition test consisted of 24 trials, including the original 12 repeated and 12 randomly generated configurations, presented in randomized order. Participants had to indicate by keyboard button press whether they had seen the displays during the course of the experiment or not. No feedback was given in the recognition task.

### Analysis

All statistical analyses were carried out using R-statistics (R Development Core Team, 2007). Experimental blocks were aggregated to four epochs, each containing five blocks. As one block contains only 12 novel and 12 repeated displays, we analyzed epochs in order to increase statistical power. Analyses of variance (ANOVAs) were performed using Type III sums of squares. For all statistical tests, the alpha level was set to 0.05.

Two data exclusion criteria were applied to the search time data. First, all incorrect responses were removed from the search time analyses because participants may not have completed search until the target was found. Second, trials in which the search time was shorter than 200 ms or larger than 3.5 standard deviations from the participants’ average search time in the remaining trials were discarded in order to remove outliers (fast guesses and extremely long searches that may unduly bias the results). This led to the rejection of 4.4% (SD = 2.7%) of invalid data for the control group, 5.0% (SD = 3.9%) for handball players, and 5.6% (SD = 4.2%) for action video game players.

## Results

### Search Times

A repeated measures ANOVA with configuration (repeated, novel), epoch (1, 4), and set size (8, 12) as within-subjects factors and group (control, handball players, action video game players) as between subjects factor was performed on search times (Figure 3).

**Figure 3:**
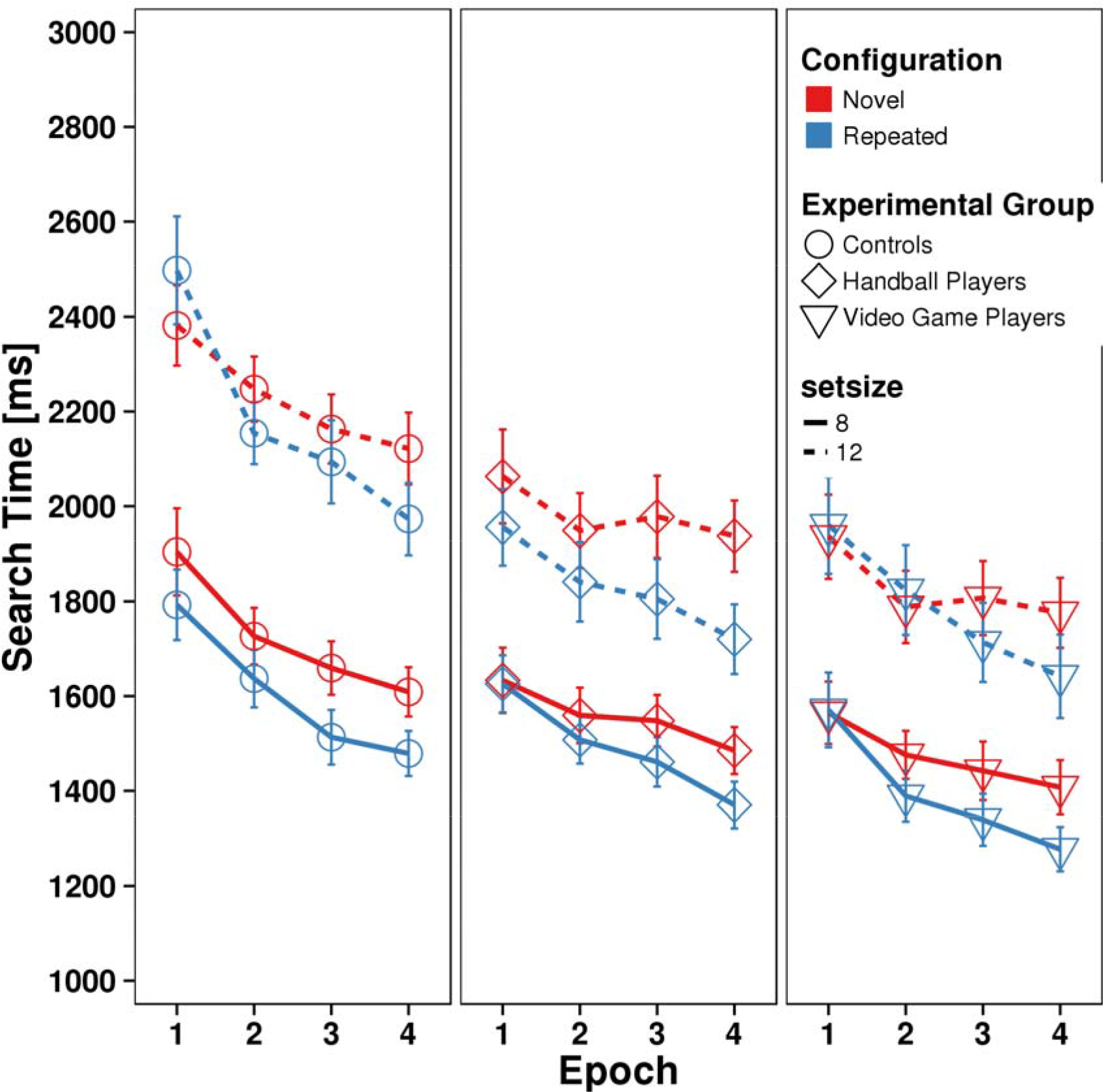
Mean search times in the visual search task. Error bars represent the standard error of means. Search times are plotted across epochs, each containing 5 search blocks, for controls (left), handball players (middle) and action video game players (right), separated for novel (red) and repeated (blue) search displays and set size eight (solid lines) and twelve (dashed lines).

**Figure 4:**
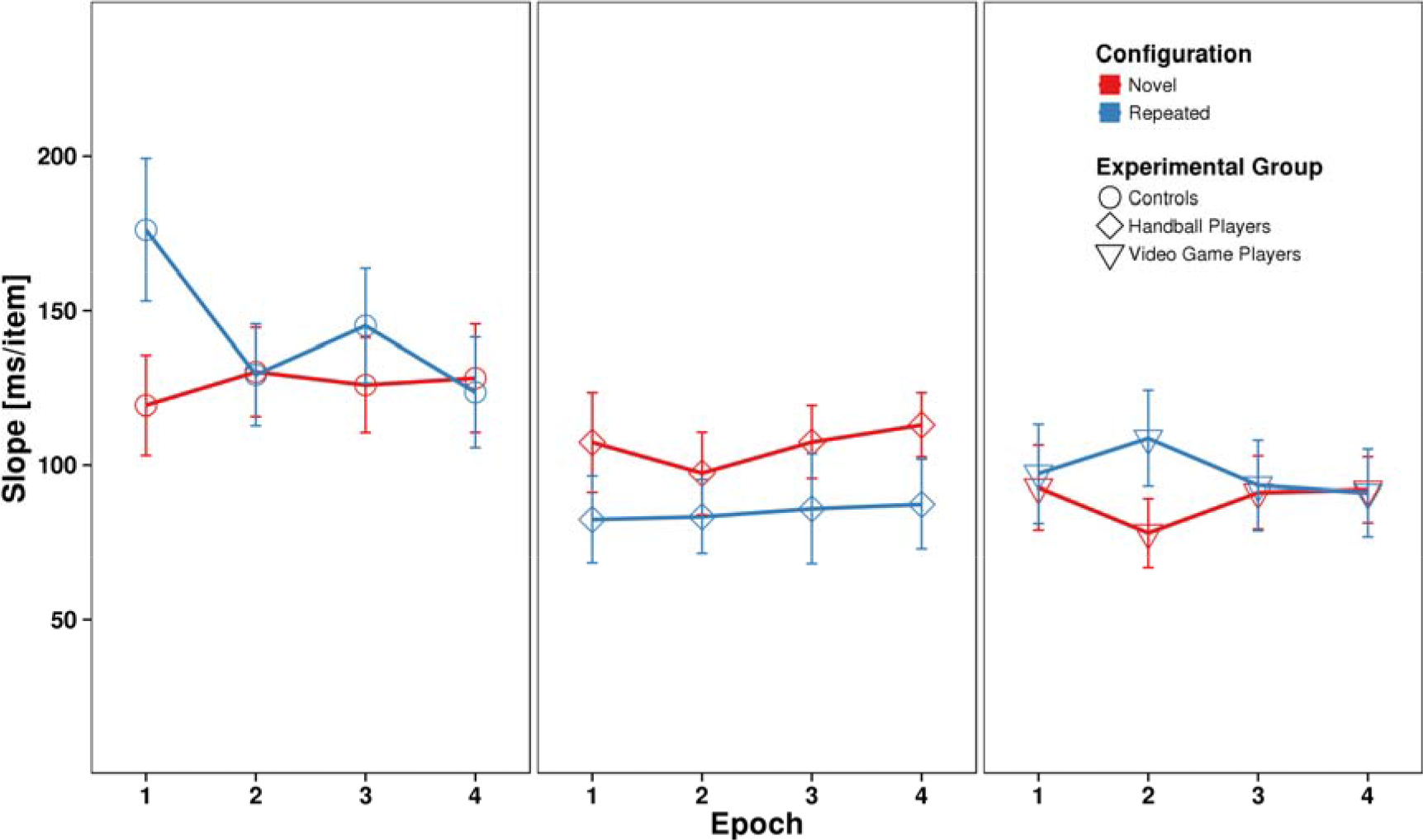
Mean search slopes of the regression line of search times. Error bars represent the standard error of means. Slopes are plotted across epochs, each containing 5 search blocks, for controls (left), handball players (middle) and action video game players (right), separated for novel (red) and repeated (blue) search displays.

**Figure 5:**
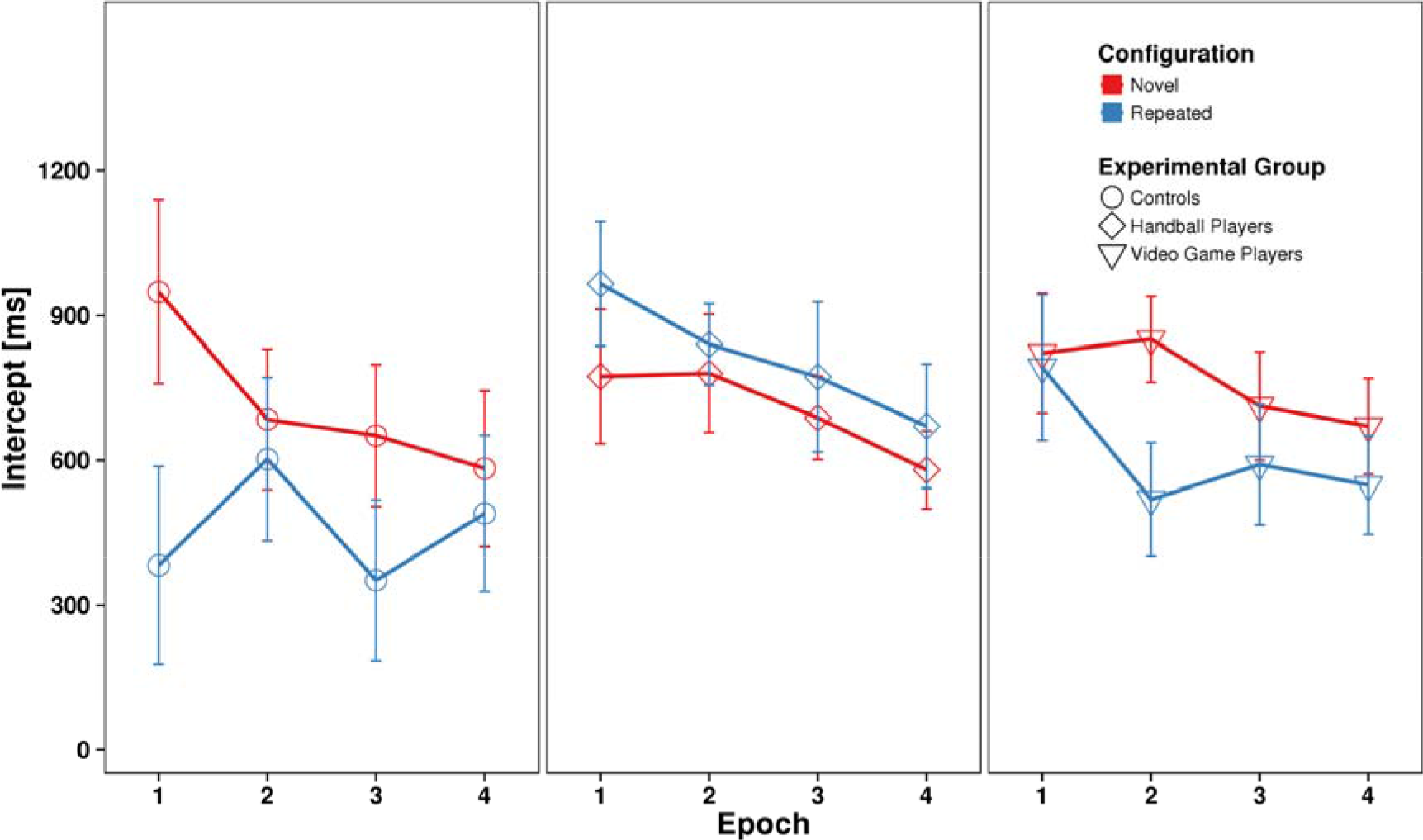
Mean intercepts of the regression line of search times. Error bars represent the standard error of means. Intercepts are plotted across epochs, each containing 5 search blocks, for controls (left), handball players (middle) and action video game players (right), separated for novel (red) and repeated (blue) search displays.

We observed a significant main effect of group [F(2,72) = 7.331, *p* < 0.05]. Post-hoc t-tests indicated that overall search speed was comparable between handball players (1808 ms) and action video game players (1773 ms; t(47) = 0.909, *p* = 0.37). However, both groups outperformed controls (1998 ms; handball players vs. control: t(48) = 2.890, *p* < 0.05; action video game players vs. control: t(48) = 3.566, *p* < 0.05). The significant main effect of epoch [F(1,72) = 97.494, *p* < 0.05] indicated general learning from the first (2100 ms) to the last epoch (1696 ms). Moreover, significant main effects of configuration [F(1,72) = 29.047, *p* < 0.05] and set size [F(1,72) = 350.173, *p* < 0.05] were found. Search times were shorter for repeated displays (1811 ms) than for novel displays (1892 ms) and for set size 8 (1557 ms) in comparison to set size 12 (2149 ms). The significant interaction of epoch x configuration [F(1,72) = 25.287, *p* < 0.05] revealed contextual cueing. Search times decreased by 318 ms in novel displays from the first to the last epoch and by 488 ms in repeated displays. Moreover, we observed significant interactions between group x epoch [F(2,72) = 3.264, *p* < 0.05] and group x set size [F(2,72) = 5.631, *p* < 0.05]. Handball and action video game players, beginning with shorter search times in Epoch 1, showed less search time reduction over epochs than controls. Larger set size slowed search much more in controls than in the other two groups. No other main effects or interactions were significant [all F <= 3.16, *p* > 0.05].

The non-significant result for the critical group x configuration x epoch interaction - indicative of contextual cueing differences across groups - could be due to either equivalence of contextual cueing scores or lack of statistical power (Dienes, 2014). To investigate this alternative further, we calculated a Bayesian repeated measures ANOVA in JASP (JASP Team, 2018, Version 0.8.2) on the group x epoch interaction of the contextual cueing effect ((RT in novel displays - RT in repeated displays) / RT in novel displays) and obtained a BF_01_ = 12.03, i.e. strong support for equivalence of contextual cueing scores across groups and epochs. Likewise, the group x epoch x set size interaction of contextual cueing scores yielded a BF_01_ = 7.28, i.e. moderate support for equivalence of contextual cueing scores.

Some of the handball players and video game players had taken part in similar experiments before (see methods). To rule out that prior experience influenced the results, in particular the group main effect, we ran an additional ANOVA, analogous to the one reported above, in which these participants were excluded (12 handball players, 9 video game players). Due to the reduced sample size, handball players and video game players were combined into one group. The ANOVA yielded a comparable pattern of results as in the main analysis, in particular a significant group main effect [F(1,53) = 5.622, *p* < 0.05]. The only differences were a non-significant group x epoch interaction [F(1,53) = 1.859, *p* > 0.05] and a significant four-way interaction [F(1,53) = 8.274, *p* < 0.05], reflecting the strong search time reduction over time for repeated displays of large set size in the control group (Figure 3), as confirmed by separate ANOVAs on search times for the control group and the experimental groups (handball and video game players) with configuration (repeated, novel), epoch (1, 4), and set size (8, 12) as within-subjects factors. In addition to significant main effects of epoch [control group: F(3,72) = 29.251, *p* < 0.05; experimental group: F(3,87) = 22.788, *p* < 0.05], configuration [control group: F(1,24) = 19.889, *p* < 0.05; experimental group: F(1,29) = 22.288, *p* < 0.05] and set size [control group: F(1,24) = 204.587, *p* < 0.05; experimental group: F(1,29) = 154.997, *p* < 0.05], in the control group, significant interactions of epoch x configuration [F(3,72) = 2.978, *p* < 0.05] and epoch x condition x set size [F(3,72) = 2.771, *p* < 0.05] were found that were absent in the experimental group [all interactions F <= 2.60, *p* > 0.05].

Independent samples t-tests (Welch t-test) on mean age between groups reveal significant differences on mean age for control group vs. handball players [t(41) = 3.143, *p* = 0.003] and for action video game players vs. handball players [t(30) = - 2.259, *p* = 0.031]. T-test on mean age between control group and action video game players were not significant [t(33) = 1.744, *p* = 0.09]. The age differences between the handball players and the other groups were due to the group of junior handball players (see participants section). To test for potential age effects on mean reaction times, we calculated an ANOVA with the within-subject factors configuration (repeated, novel) and epoch (1, 4) and the between-subject factor group (junior handball players, adult handball players). While significant main effects of epoch [F(1,23) = 16.553, *p* < 0.05], configuration [F(1,23) = 13.704, *p* < 0.05], set size [F(1,23) = 95.177, *p* < 0.05] and a significant interaction of epoch x configuration [F(1,23) = 4.293, *p* < 0.05] replicated our analyses above, importantly, no significant main effect of group or interaction effects involving the group factor were observed [all F <= 3.11, *p* > 0.05]. Thus, we have no evidence for an age effect.

It might be argued that the lower search times that we observed in Epoch 1 for the athletes and video game players relative to the control group were potentially due to fast learning in Epoch 1 in the former two groups. To analyze this hypothesis, we ran an additional ANOVA on the Epoch 1 search times with configuration (repeated, novel), block (1 - 5), and set size (8, 12) as within-subjects factors and group (control, handball players, action video game players) as between subjects factor. We observed significant main effects of group [F(2,72) = 7.474, *p* < 0.05], block [F(4,288) = 7.870, *p* < 0.05], set size [F(1,72) = 299.923, *p* < 0.05], and a significant group x set size interaction [F(2,72) = 7.891, *p* < 0.05]. The group x condition x set size interaction narrowly missed significance [F(1,72) = 3.077, *p* = 0.052]. All other effects were not significant [all F <= 1.490, all *p* > 0.16]. Thus, importantly, we observed no significant interactions involving group x block that might indicate different learning rates of the groups in Epoch 1.

We further investigated potential effects of sex on search time with an ANOVA with sex as between-subject factor and configuration, epoch and set size as within-subject factors. This analysis yielded no significant main effect of sex [F(1,73) = 0.138, *p* = 0.71] and no significant interactions involving sex [all F <= 2.44, *p* > 0.12.].

For further analysis of the contribution of attentional and post-selective processes contained in the search times, we investigated the slopes and intercepts of the search time x set size function (Figures 4 and 5). A repeated measures analysis of variance (ANOVA) with configuration (repeated, novel), and epoch (1, 4) as within-subject factors and group (control, handball, video) as between subjects factor was performed separately on slope and intercept data.

#### Slopes

As an indicator for search efficiency, we calculated the slopes of the search time x set size regression lines. We observed a main effect of group [F(2,72) = 6.502, *p* < 0.05] on slopes. No other main effects or interactions were significant (Table 1). Post-hoc t-tests revealed that mean search times per item were higher for controls (135 ms / display item) than for handball players (96 ms, t(47) = 2.661, *p* < 0.05) or action video game players (93 ms; t(47) = 2.990, *p* < 0.05), but did not differ between handball and action video game players [t(48) = 0.317, *p* = 0.75].

**Table 1:** Statistical results of the slope and intercept analyses.

| Effect | Measure |  |  |  |  |  |
| --- | --- | --- | --- | --- | --- | --- |
|  | Slopes |  |  | Intercepts |  |  |
| | F | p | $\eta^2$ | F | P | $\eta^2$ |
| Group | 6.50 | **.003** | .062 | 1.36 | 0.26 | .011 |
| Condition | 0.04 | 0.85 | .000 | 1.39 | 0.24 | .005 |
| Epoch | 0.65 | 0.58 | .002 | 4.13 | **.007** | .012 |
| Group:Condition | 1.75 | 0.18 | .013 | 1.60 | 0.21 | .013 |
| Group:Epoch | 0.63 | 0.70 | .003 | 0.27 | 0.95 | .002 |
| Condition:Epoch | 1.21 | 0.31 | .003 | 0.25 | 0.86 | .001 |
| Group:Condition:Epoch | 1.70 | 0.12 | .009 | 1.66 | 0.13 | .009 |

Again, we repeated the analysis excluding participants with prior experience in contextual cueing experiments. This ANOVA confirmed the significant group main effect [F(1,53) = 4.531, *p* < 0.05]. The only other significance was observed for the three-way interaction [F(1,53) = 8.274, *p* < 0.05], reflecting the large decrease of the search slope over time for repeated displays in the control group.

#### Intercepts

An analogous repeated measures ANOVA as for slopes was calculated on the intercepts. The intercept analysis revealed a significant main effect of epoch [F(1,72) = 9.207, *p* < 0.05], indicating a reduction of intercepts over the course of learning. No other main effects or interactions were significant (Table 1). An ANOVA excluding participants with prior experience in contextual cueing confirmed the significant main effect of epoch [F(1,53) = 5.019), *p* < 0.05]. The only other significant effect was observed for the three-way interaction, reflecting the increase of the intercept from Epoch 1 - 4 for repeated displays in the control group.

All statistical results of the between-group analyses of contextual cueing for slope and intercept data are reported in Table 1.

### Accuracy

We observed very high average accuracies for all groups (98.1% for the control group, 97.1% for the handball players and 97.1% for the action video game players; Table 2). An ANOVA on the logit-transformed accuracy data with the between-subjects factor group (control, handball, video) and the within-subject factor configuration (repeated vs. novel) yielded no significant effects [all F<= 2.226, *p* > 0.05]. To test for equivalence, we calculated an analogous Bayesian ANOVA on the logit-transformed accuracies. It yielded a BF_01_ = 3.71, i.e. moderate support for equivalence of accuracies between groups (main effect). The group x configuration interaction yielded a BF_01_ = 6.37, i.e. moderate support for equivalence of group effects across configurations.

**Table 2:**
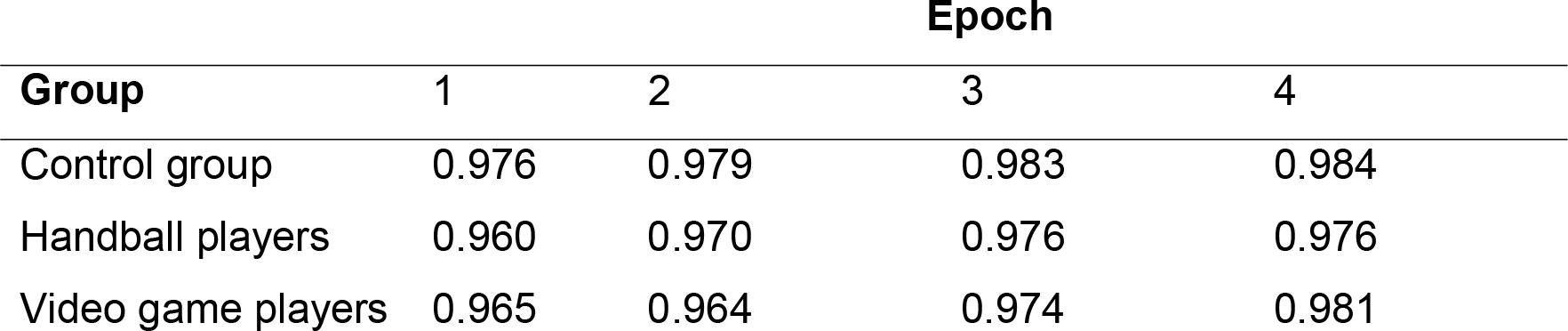
Accuracy data by group and epoch.

| Group | Epoch |  |  |  |
| --- | --- | --- | --- | --- |
|  | 1 | 2 | 3 | 4 |
| Control group | 0.976 | 0.979 | 0.983 | 0.984 |
| Handball players | 0.960 | 0.970 | 0.976 | 0.976 |
| Video game players | 0.965 | 0.964 | 0.974 | 0.981 |

Again, we repeated the analysis excluding participants with prior experience in contextual cueing experiments. High accuracies (96.5%) for the participants without experience were observed. The ANOVA on accuracy for the subgroup without prior experience confirmed the non-significant effects obtained for the whole group [all F<= 2.455, *p* > 0.05].

### Recognition Task

The control group obtained a mean recognition accuracy of 58.7% (SD = 10.3%). The mean hit rate of 58.7% (SD = 17.6%) and the mean false alarm rate of 41.3% (SD = 13.5%) differed significantly [t(24) = 4.1903, *p* < 0.05]. However, the correlation between the standardized contextual cueing effect and the recognition accuracy was not significant (r = 0.095, *p* > 0.05).

Mean recognition accuracy for the handball players was 56.8% (SD = 13.2%). Their mean hit rate of 56.7% (SD = 10.8%) and the mean false alarm rate of 43.0% (SD = 21.3%) differed significantly [t(24) = 2.594, *p* < 0.05]. The normalized contextual cueing effect, again, did not correlate with recognition accuracy (r = - 0.047, *p* > 0.05).

Action video game players reached a mean recognition accuracy of 52.2% (SD = 11.6%). Hits (M = 49.0%, SD = 18.7%) and false alarms (M = 44.7% (SD = 11.8%) were not significantly different [t(24) = 0.933, *p* > 0.05], yielding no indication of explicit learning. Again, recognition accuracy did not correlate with the normalized contextual cueing effect (r = −0.615, *p* > 0.05).

## Discussion

We investigated if high-level amateur team sport players and action video game players show a superior visual search performance in arbitrary, non-sport specific search configurations. In particular, we investigated if visual search performance in these groups benefits from superior search guidance and/or context learning abilities.

Participants searched for a T-shape among L-shaped distractors and had to indicate the orientation of the T-shape by an alternative forced choice button press. Half of the search configurations were repeated across blocks so that the configurations could be learned and used to guide search for the target location. Importantly, the orientation of the target was always random so that no specific response could be associated with any repeated display.

We found that both handball players and action video game players searched faster than controls. This search time advantage was analyzed further in that we calculated search slopes, i.e. search time increase per display item, as a measure of search efficiency. Search times per display item were shallower in both handball players and action video game players, indicating that the members of both groups needed less time to analyze the contents of a search display than controls. This is in agreement with previous work that reported shorter search times per item in action video game players than controls in inefficient search tasks such as the one used here (Hubert-Wallander et al, 2011; Wu, & Spence, 2013). While slopes are defined by search duration per item, this should not be taken literally. Shallower slopes could mean faster sequential processing of the search display, but it could also mean faster parallel processing of the display, for instance using a larger attentional focus. We cannot distinguish between this alternative on the basis of the current data. However, this question could be addressed in future work using eye-tracking or reaction time modeling (e.g. Müller-Plath, & Pollmann, 2003). We also note that our interpretation depends on the assumption of two - selective and postselective - processing stages but other models are of course possible and may lead to different interpretations (Kristjansson, 2015),

In contrast, no difference between groups was observed regarding the intercepts of the search time regression curve with the y-axis. The intercept indicates fixed time “costs” that are independent of display size and may arise due to response preparation or execution. In the context of visual search, slopes and intercepts are usually interpreted as indicators of attentive respectively post-selective processes (e.g. Kunar et al., 2007, but see criticism by Kristjansson, 2015). In this framework, the superior search performance of both athletes and video game players could be attributed to superior attentive processing. This pattern was confirmed when athletes with prior experimental experience were excluded from the analysis.

In addition, the comparison of novel and repeated display configurations enabled us to investigate spatial configuration learning. Although context learning was incidental, i.e. participants did not know about display repetition, search times decreased more in repeated than novel displays over the course of the experiment. Thus, all groups showed incidental learning of repeated displays, in line with many previous reports on this contextual cueing effect (Chun, 2000). However, handball players and action video game players did not differ from controls (nor from each other) in the amount of search facilitation in repeated configurations. Thus, our data yield no evidence for superior learning of arbitrary scenes.

We could rule out effects of age and prior experience with psychological experiments as factors of influence. As in previous studies, we had a sex imbalance between groups, mainly due to the male dominance among the action video game players. Sex, however, did not significantly influence search times. Handball players and controls selected repeatedly presented displays with abovechance probability, indicating at least partial explicit memory. However, recognition accuracy did not correlate with the size of the contextual cueing effect (the search time reduction in repeated displays). Thus, we have no indication that contextual cueing was due to top-down controlled search based on explicit knowledge of the target location in repeated displays.

It should be noted that the improved search performance of the handball and action video game players was present from the beginning of the experiment and did not develop further over the course of the experiment. In fact, the control group showed a stronger search time reduction than the two experimental groups. Thus, the handball and action video game players already started with superior attentional capabilities and did not develop them with repeated stimulus exposure, in contrast to what has been reported in very demanding psychophysical tasks (Bejjanki et al., 2014).

Several caveats should be considered regarding superior performance of athletes or video-game players. In the present study, we needed to rely on open recruiting of semi-professional handball players, because they would not occur frequently enough in a random sample. Selection of special groups, however, may go along with the motivation to perform well (Boot et al., 2011), particularly because reports of superior performance, particularly of video game players, have been published in the general media. However, the accuracy data yielded no indication of a speed-accuracy trade-off. Nevertheless, we cannot completely rule out that search speed was more affected by motivation than contextual cueing, perhaps because the former is an evident goal of a search task, whereas display repetition is not announced and often not consciously perceived, or only perceived late during the task.

Moreover, we do not know if the superior search performance of the handball players and action video game players is a training or a selection effect. Longitudinal studies would be needed to investigate if handball or action video game training leads to improved search performance. Alternatively, it may be that persons with superior attentional processing skills are more likely to become successful handball or action video game players (Kristjansson, 2013).

Furthermore, our findings do not imply that handball players may not have better memory-guided search in realistic handball scenes. In fact, across many studies, sport-specific displays, stimuli, and processing requirements were more likely to lead to expert-novice differences (Abernethy, 1987, 1988; Mann et al., 2007). Our results, however, do not support the view that handball players or video game players have better memory-guided search outside of their domains of expertise.

To conclude, we replicated previous reports of faster visual search in athletes and in action video game players. In addition, we observed that the superior search speed was due to faster attentional processing, whereas response-related processes did not differ from the control group. In contrast, handball players or action video game players showed no better-than-normal attentional guidance by learned spatial contexts.

### Compliance with ethical standards

All procedures performed in studies involving human participants were in accordance with the ethical standards of the institutional and/or national research committee and with the 1964 Helsinki declaration and its later amendments or comparable ethical standards. Informed consent was obtained from all individual participants included in the study.

## Acknowledgements

This work was supported by the Deutsche Forschungsgemeinschaft (PO548/14-1 and SFB779-A4).

## Author contributions

A.S. designed the study, wrote the experimental code, acquired and analyzed the data and wrote the manuscript. F.G. wrote the experimental code and analyzed the data. F.S analyzed the data. S.P. designed the study and wrote the manuscript.

## Additional information

The authors declare no competing financial interests.

